# Changes in movement characteristics in response to private and social information acquisition of socially foraging fish

**DOI:** 10.1101/2021.07.30.454431

**Authors:** Geoffrey P. F. Mazué, Maxim W. D. Adams, Frank Seebacher, Ashley J. W. Ward

**Author notes:** Correspondence should be addressed to G. P. F. Mazué. **AUTHORS’ CONTRIBUTIONS** G.P.F.M. and A.J.W.W. designed the experiment; G.P.F.M. performed the experiment; G.P.F.M. and M.W.D.A. scored the video recordings and performed the tracking; G.P.F.M. analysed the data with inputs from F.S. and A.J.W.W.; G.P.F.M. wrote the manuscript. All authors edited the manuscript. **DATA AVAILABILITY STATEMENT** Data will be deposited in the Dryad Digital Repository upon acceptance of the manuscript.

## Abstract

To overcome the cost of competition resulting from foraging in a group, individuals may balance their use of private (i.e., acquired from personal sampling) and social (i.e., acquired by watching other individuals) information to adjust their foraging strategy accordingly. Reliability of private information about environmental characteristics, such as the spatial distribution of prey, is thus likely to affect individual movement and social interactions during foraging. Our aim was to investigate how movement characteristics of foraging individuals changed as they acquired reliable private information about the spatial occurrence of prey in a foraging context. We allowed guppies (*Poecilia reticulata*) to develop the reliability of their private knowledge about prey spatial occurrence by repeatedly testing shoals in a foraging task under three experimental distributions of prey: 1) aggregated prey forming three patches located in fixed locations, 2) scattered distribution of prey with random locations, or 3) no prey (used as control). We then applied tracking methods to obtain individual time series of spatial coordinates from which we computed a suite of movement variables reflecting search effort, social proximity and locomotion characteristics during foraging, to examine changes occurring over repeated trials and to investigate which best explained foraging success. We show that foraging shoals became more efficient at finding and consuming food over the first three days by increasing their time spent active. Over time, individuals foraging on either scattered or aggregated prey travelled greater distances, showed an increasing distance to their closest neighbour and became slightly more stochastic in their acceleration profile, compared to control individuals. We found that behaviour changed as private information increased over time, with a clear behavioural shift occurring on the third testing day. Social proximity was the major predictor of foraging success in the absence of prior foraging information, while search effort became the most important predictors of foraging success as information increased. In conclusion, we show that individual movement patterns changed as they acquired private information. Contrary to our predictions, the spatial distribution of prey did not affect any of the movement variables of interest. Our results emphasise the importance of information in shaping movement behaviour in animals.

## INTRODUCTION

Individuals benefit from foraging socially through diverse mechanisms such as local enhancement (e.g., cues from successful foragers provide information about food location) or social facilitation (e.g., initiating foraging quicker in the presence of foraging conspecifics), which allow an increase in food detection, information dissemination and foraging success (Galef & Giraldeau, 2001; Giraldeau & Beauchamp, 1999; Giraldeau & Caraco, 2018; Grand & Dill, 1999; Pitcher et al., 1982; Snijders et al., 2021; Ward & Zahavi, 1973). In a social foraging scenario, two types of information are available: ‘private’ information, which individuals acquire from their own experience within the environment such as the recent observation of prey in the vicinity or the memory of a location associated with recent successful foraging events (Valone, 1991); and ‘social’ information which is acquired from watching other foragers and gauging their success and the level of competition at a patch (Coolen et al., 2003; Templeton, & Giraldeau, 1995; Valone, 1989). At any moment, an individual faces the choice of continuing to search for food on its own or joining other foraging conspecifics. Such a decision therefore results from balancing private and social information to reduce uncertainty about the environment and so maximise foraging success (Templeton & Giraldeau, 1996). Social information is typically ubiquitous in social foraging and it is assumed to be less costly to acquire and use in terms of time and energy than private information. Yet, relying excessively on social information can lead individuals to make maladaptive decisions (Giraldeau et al., 2002; Laland & Williams, 1998), and foraging within close social proximity can generate non-negligible costs of competition (Cresswell, 1998). Additionally, individual nine-spined sticklebacks (*Pungitius pungitius*) tended to disregard social information when making foraging decisions if their private information was reliable and up-to-date (Van Bergen et al., 2004). Similarly, guppies (*Poecilia reticulata*) relied on private knowledge and preferentially foraged on patches they had visited before, rather than joining an alternative patch occupied by conspecifics (Kendal et al., 2004). Therefore, individuals might benefit from relying preferentially on private information when its acquisition is recent and when its cost is low (i.e., time and energy allocated to private sampling, risk of predation).

In most complex natural environments, food availability may vary both over time and in its spatial distribution (Riotte-Lambert & Matthiopoulos, 2020), which necessitates continuous gathering and processing of information in order to adjust foraging behaviour (Higginson et al., 2012). In the absence of private information about the spatial or temporal occurrence of resources in the environment, social information is more likely to be relied on (Harel et al., 2017; Lewis et al., 2013; Reebs, 2000). As they sample the environment, individuals increase their private information about food distribution. Experience improves a forager’s ability to predict the outcome of future sampling at a given location and/or time. Private information that promotes successful foraging may then be considered as reliable information. Such reliability is thus likely to affect the decisions that individuals make, such as how they move in the environment and how they interact with other foragers (Riotte-Lambert & Matthiopoulos, 2020), and it is likely to be affected by the distribution patterns of the resources. For example, foraging on unpredictable patchy resources may promote grouping behaviour and the use of social information to maximize chances of discovery and minimize the latency to arrive at a newly-discovered food patch (Beauchamp & Ruxton, 2014; Ioannou et al., 2011). However, in the presence of private information about patch location, foraging asocially may reduce costs of competition within a foraging patch. Alternatively, when food discoveries lead to relatively small food intakes, as is the case when resources are distributed more evenly across the environment, individuals should ignore social information to rely mostly on private information and effort search (Humphries et al., 2012).

In this study, we investigated the effect of reinforced private information on movement characteristics of individuals foraging on different spatial distributions of prey. We analysed high resolution trajectory data collected from shoals of guppies (*Poecilia reticulata*) that were allowed to forage repeatedly on prey that were either aggregated in patches at constant locations or scattered at random locations in the testing arena. First, we examined the changes occurring in a suite of variables describing different aspects of individual foraging behaviour including distance travelled as a measure of search effort, distance to nearest neighbour as a measure of social proximity, and acceleration entropy as a measure of stochasticity in locomotion decisions. Patterns observed in both foraging treatments were compared to those of control shoals tested in absence of food. Then we estimated relative importance of these variables at explaining individual foraging success in the two foraging treatments. We hypothesised that (*i*) in the absence of prior information about the prey distribution and competition, individuals initially forage as a group to enhance their exposure to social information, regardless of the prey distribution; and (*ii*) changes of behaviour occur as private information about spatial occurrence of prey and perceived level of competition increases, before reaching a phase of fixation when individuals’ knowledge of the foraging conditions has become highly reliable (i.e., reinforced). A corollary from (*ii*) is that (*iii*) the spatial distribution of prey should affect behavioural adjustments over time. We predicted the development of an asocial foraging strategy in individuals exposed to scattered prey characterized by increasing search effort, social proximity and stochasticity in locomotion over days, and the development of a more social strategy in individuals exposed to aggregated prey characterized by a pronounce social proximity (i.e., short distances to neighbour) and a goal-oriented movement (i.e., small distance travelled, with less stochasticity in locomotion).

## MATERIALS AND METHODS

### Experimental subjects

We used 140 female domestic guppies (*Poecilia reticulata*) as experimental subjects (obtained from Pisces Aquatics, Kenmore, QLD, Australia). Individuals had a body length of 33.7 ± 2.8 mm (standard length, mean ± sd). Only females were used in this experiment as previous research showed that sex affects movement, food discovery and information spreading in foraging shoals of guppies; with females having greater success at discovering foraging patches than males (both under laboratory and natural conditions) and enhanced spreading of foraging information (i.e., route to access the foraging patch) in groups of females than groups of males (Croft et al., 2003; Reader & Laland, 2000; Snijders et al., 2019). Individuals were split between two large 180L white plastic tanks placed in a temperature and photoperiod (12h Light: 12h Dark cycle) controlled room for 22 days before commencement of the experiment. Holding tanks were filled with dechlorinated aged water maintained at a constant temperature of 23°C and furnished with air powered sponge filters and plastic plants. Fish were fed tropical fish flakes (SERA Vipan, Heinsberg, Germany) twice daily during the first 12 days, and then received flakes in the morning and defrosted bloodworms (*Chironomidae*, Ocean Nutrition) in the afternoon for the remaining 10 days in order to ensure the fish were familiar with the prey later used in the experiment (Godin, 1978). The day before the commencement of the experiment, fish were gently netted from their holding tank and distributed across 12 experimental groups of four individuals. Each fish within a group had a unique coloration patterns on their abdomen and their caudal fin, enabling individual identification in video recordings. Each experimental group was photographed from above in a small container containing a ruler placed in shallow water to enable the measurement of each individual’s standard length from the photograph using ImageJ. Each experimental group was placed in an opaque green plastic tank (33 cm L x 24 cm W x 18 cm H, 14L) containing an air powered sponge filter, and two plastic plants, and filled with dechlorinated aged water maintained at a constant temperature of 23°C and kept under the same light cycle as before. Two blocks of 12 groups and a third block of 10 groups were tested within eight consecutive weeks (October-December 2019), and treatments were balanced within each block.

### Experimental design

We used two identical experimental arenas made of opaque white Perspex (inner dimensions 118 cm L x 30 cm W x 20 cm H). The bottom of the arenas was covered with a 8 mm deep white plastic grid with each square of the grid measuring 15 mm x 15 mm (referred to as ‘foraging pits’). Each of the 1200 foraging pits contained either a single prey item or none, and ensured that prey were held in assigned locations and did not drift in currents generated by nearby swimming fish. We used a standardized quantity (n=12) of defrosted bloodworms (*Chironomidae*, Ocean Nutrition, Essen, Belgium) as prey items, that were distributed across the grid according to one of the three experimental treatments: 1) the ‘aggregated’ treatment (n=13 groups) consisting of 12 bloodworms divided into three aggregations of 4 adjacent foraging pits, at fixed locations across the grid; 2) the ‘scattered’ treatment (n=12 groups) consisting of 12 bloodworms scattered across the arena with a pattern of distribution changing daily, obtained by haphazardly scattering bloodworms while avoiding any aggregation between two worms; and 3) the ‘control’ treatment (n=10 groups) consisting of an empty foraging grid containing no prey items (Figure 1). The arena was filled with 7 cm of dechlorinated aged water, leaving 6.2 cm of water column for the fish to navigate above the grid. Such a low water depth reduced the ability of the fish to see multiple foraging pits while swimming, forcing them to discover which foraging pits contained a prey item by exploring the entire arena. A filtered solution from 10 crushed bloodworms was added to the arena each day before the trials commenced to standardize the amount of foraging cues in the arena for the first groups to be tested on the day. Three sandstone cylinders (diameter 5.8 cm and height 8 cm) were placed at fixed locations (across replicates and treatments) of the grid to create environmental enrichment and break-up the line of sight of foraging individuals. A square container made of transparent plastic (15 cm side lengths) was used as a starting box (Figure 1). The arenas were illuminated overhead with fluorescent tubes and were surrounded by white plastic sheets to minimize external stimuli.

**Figure 1.**
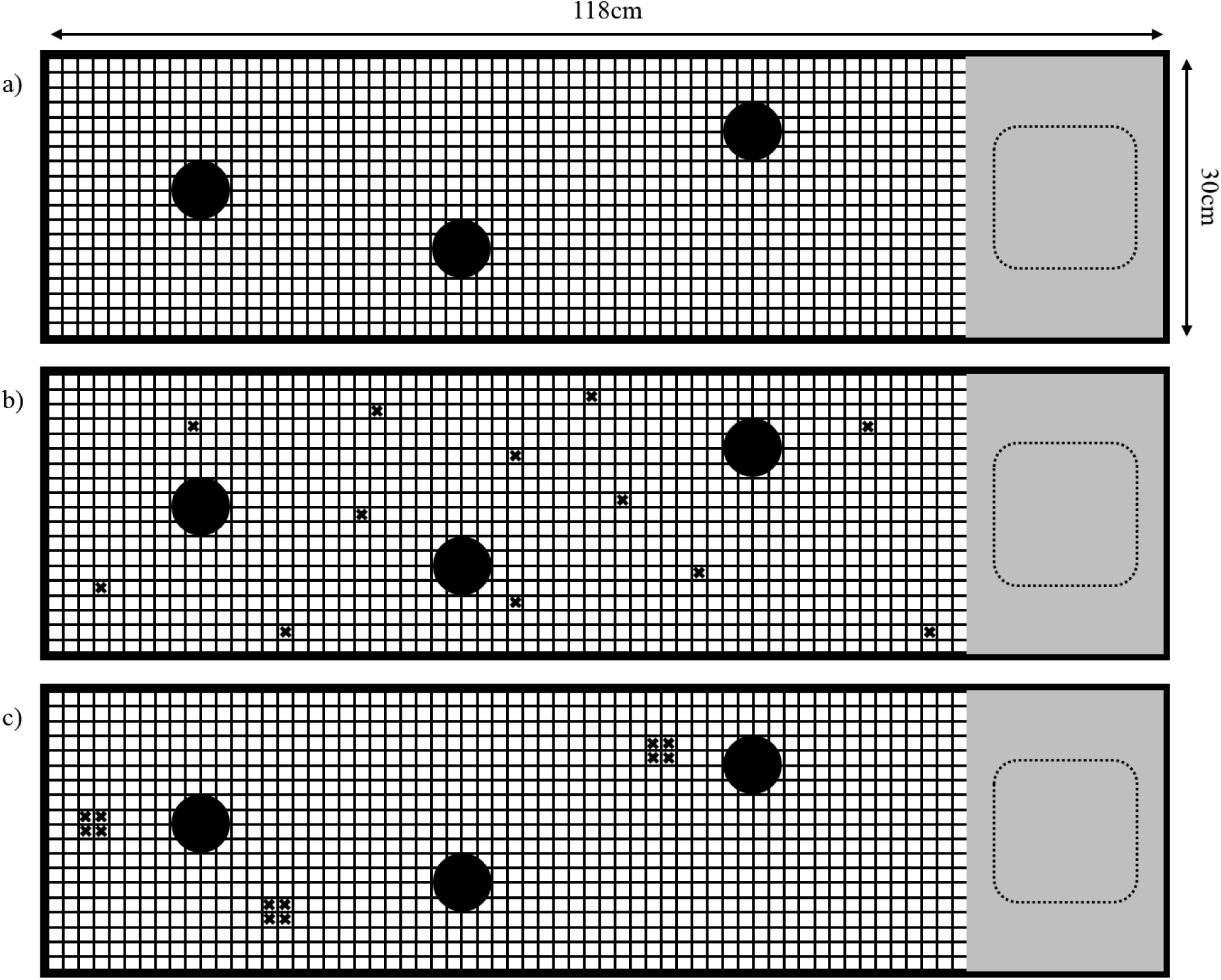
Top view of the experimental arena with the spatial distributions of prey used as experimental treatments: a) control, b) scattered, and c) aggregated. The dashed-line square symbolises the starting box used to release the fish, set on a platform symbolized by the grey area. Sandstone cylinders are symbolized by black disks. Prey items (i.e., bloodworms) are symbolized by small black crosses.

Experimental groups were randomly assigned to one of the three treatments on the day prior to the commencement of the experiment and were tested daily for 10 consecutive days. Testing order and arenas were randomized daily to prevent any effect on foraging behaviour. Tested groups were gently netted out of their holding tank to be placed inside the starting box. After 2 minutes of acclimation, one edge of the starting box was raised remotely using a pulley system, freeing the fish to explore the arena. All trials were filmed for 25 min at 30 frames per second and at a resolution of 1280×720 pixels, using a Canon G1X camera suspended above the arena. After completion of a trial, each group was returned to its holding tank. To standardize the food intake, all groups assigned to the control treatment received 12 bloodworms in their holding tank. All groups were offered tropical fish flakes (SERA Vipan) at the end of the day.

### Data extraction and processing

From the video recordings, we recorded the latency (in frames, measured from the frame when the starting box was raised) for each bloodworm to be eaten and the identity of the fish that consumed it. We then quantified the time (in seconds) for the group to consume the 12 bloodworms (later referred as ‘depletion time’). Groups that failed to find and consume all bloodworm within that a maximal time of 25 min were given a score of 1500sec (3 groups on day 1). Early inspection of the data revealed that the depletion time decreased significantly over the first 4 days to reach a plateau at day 5 (Figure 2). Subsequently, further data extraction and analyses were performed solely on the days 1, 3, 5 and 7 of the experiment.

**Figure 2.**
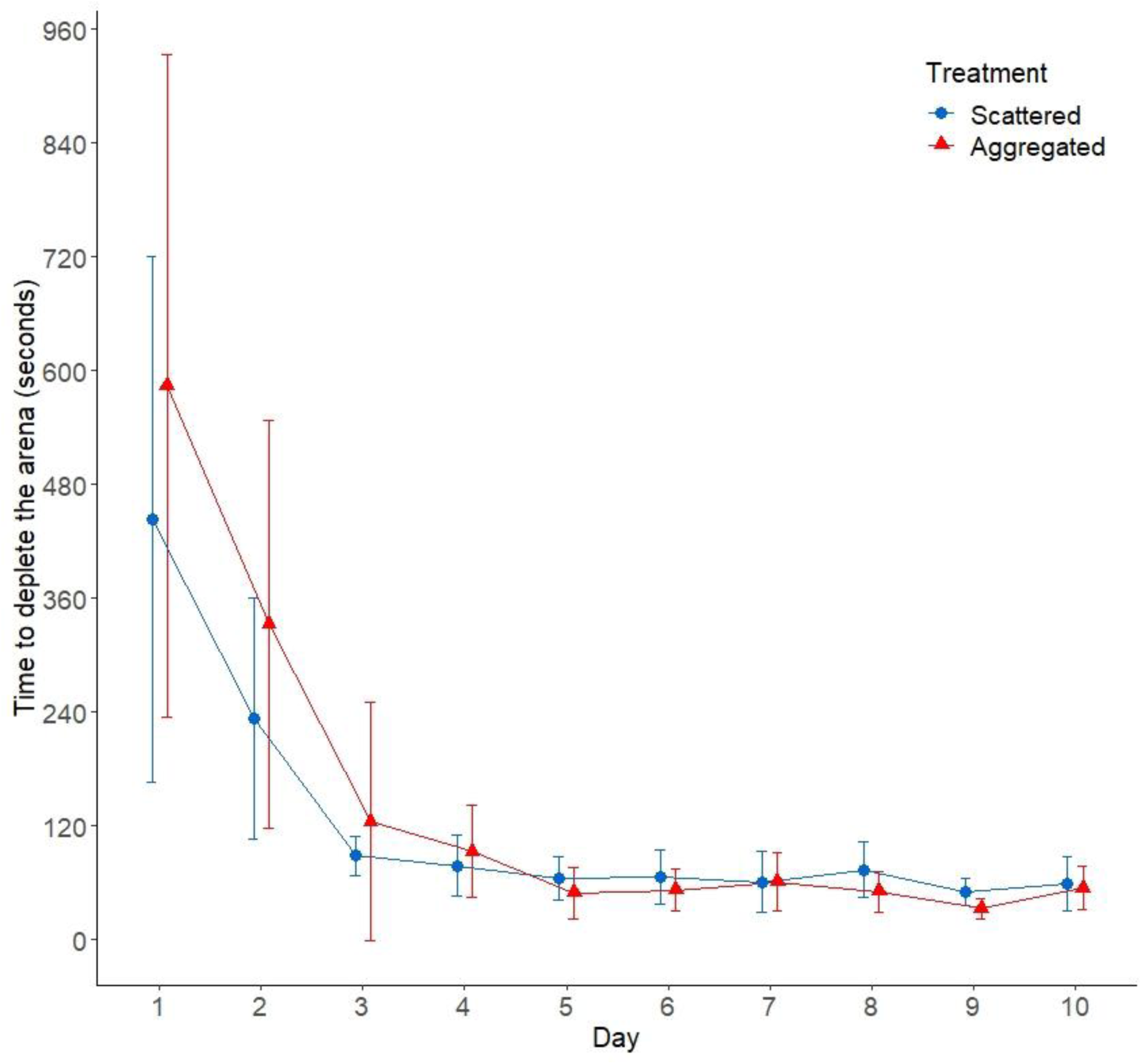
Visualisation of collective foraging efficiency. Time elapsed until the depletion of the arena (i.e., when the last prey was consumed) as a function of treatment and for the 10 consecutive days (seconds, mean ± 95%CI). Only ‘scattered’ (blue) and ‘aggregated’ (red) treatments are displayed because there was no prey in the ‘control’ treatment.

To track individual movement, video recordings from the aggregated and scattered treatments were cropped from the ‘starting frame’, defined as the frame at which the first bloodworm was picked by a fish (as a standardization method), onwards to reach a total of 1800 frames (i.e., 60 sec). This period was chosen because most of bloodworms were consumed during the first minute. Videos from the control treatment, were instead cropped to 1800 frames from a starting frame equal to the averaged value of the starting frames for the aggregated and scattered videos of the same day. Cropped video files were then converted to .avi using VirtualDub (version 1.10.4) and tracked using the CTRAX automated tracking software (Caltech Ethonomics Project, The Caltech Multiple Fly Tracker, Version 0.5.18) (Branson et al., 2009). The time series of (x, y) coordinates produced for each individual were then screened for any abnormalities and manually corrected using the FixErrors GUI in MATLAB. Pixel coordinates were converted to mm using a conversion ratio calculated by measuring the distance between two points of known distance on a video frame. Within each experimental group for each daily video recording, the four time-series of coordinates (of length 1800 frames each), were matched to the identity of the fish based on the coloration patterns of their abdomen and caudal fins.

From the tracking data we performed different calculations to characterize three distinct aspects of individual movement in a foraging context and to describe grouping patterns observed at the level of the group (see SI). First, we calculated each individuals’ speed (standardized in body length per second) at each time step. Within this time series for each individual, we identified all bouts (defined as sequences of consecutive frames) for which it was moving by using an arbitrary criterion of speed greater than 0.5 BL.s^-1^ (Jolles et al., 2020). Additionally, we calculated the distance between each pair of fish (in averaged body length unit) at each time step, from which we extracted the distance of each fish to its nearest neighbour (further referred to as distance to NN, or NND) at each frame. Subsequently, we quantified the cumulative distance (in cm) travelled by each individual over all frames for which it was identified to be moving, as a measure of search effort. The average distance to NN of each individual, calculated as the median distance to NN over all frames for which it was identified to be moving, was used as a measure of social proximity while foraging. Additionally, we computed the entropy values for the time series of acceleration for each bout of at least 15 consecutive frames (i.e., half a second) during which a fish was identified as moving (a threshold determined to be highly conservative, as it contained over 80% of the frames spent moving within each day), using the ‘pracma’ package in R (Borchers, 2019). Approximate entropy estimates the amount of regularity and unpredictability of fluctuations in a time series, enabling us to determine whether individuals’ locomotion was more deterministic (low values of approximate entropy) or more stochastic (i.e., less predictable, high values of approximate entropy) while exploring and searching for food in the arena (Pincus, 1991). The median value of all bouts’ entropy values was used as an overall measure of unpredictability in acceleration patterns of a foraging fish.

Individual foraging success, defined as the number of prey consumed, was scored by giving a +2 score for each bloodworm that was consumed in its entirety during the 60s of the tracked video segments. Because of a non-negligible number of kleptoparasitism events, a score of +1 was given to both individuals that were involved in the consumption of a single bloodworm (e.g., stealing or dropping/picking up a half-consumed prey).

### Statistical analysis

Sample sizes presented until day 5 included 13 shoals (52 fish) in the aggregated treatment, 12 shoals (48 fish) in the scattered treatment and 10 shoals (40 fish) in the control treatment; but only 12 and 11 shoals (48 and 44 fish) in aggregated and scattered treatments (respectively) on day 7 due to the exclusion of two groups from the experiment caused by the loss of two individuals. Our general approach was to analyse the dependent variable using models fitted according to their initial distribution, and no transformation was applied on any of the dependent variables.

#### Analysis of depletion time

We first analysed the depletion time using a Generalized Linear Mixed Effects model (GLMM) fitted with a negative binomial error distribution. The model included day and treatment (with the exclusion of the control level as there were no prey in the arena) as fixed effects, and shoal identity as random factor. Attempts to simplify the models were made by comparing the models with and without interaction terms based on their AIC values. The significance of the terms kept in the reduced models were assessed using type III ANOVAs. Model assumptions were verified through visual inspection of the Q-Q plot of the residuals and by plotting the predicted residual against the observed residuals.

#### Analysis of movement characteristics

We analysed the effect of repeated testing (i.e., day) and prey distribution (i.e., treatment), and their interaction, on each of the three dependent variables, using GLMMs fitted with a Gaussian or Gamma error distribution depending on the estimated distribution of the dependent variables analysed. The model included treatment and day and their interaction as fixed effects, and fish identity nested within the shoal identity as random effects. Attempts to simply the models, assessment of terms’ significance, and assumption verifications were performed as described previously.

#### Analysis of the foraging success

We used a model averaging approach to assess the relative effect of our variables of interest, *viz*. total distance travelled, average distance to NN, treatment and their pairwise interactions on individual foraging success. Acceleration entropy was excluded from the analyses as it showed signs of collinearity with distance travelled (see Table S1). We calculated estimates of model-averaged coefficients (with their 95%CIs) from a set of candidate models generated from reducing an initial model containing all terms as fixed effects, as well as shoal identity used as a random term (Burnham & Anderson, 2002; Grueber et al., 2011). Foraging success was thus analysed as a response variable in a generalized mixed effect model (GLMM) fitted with Poisson error distribution. A set of 64 candidate models was generated from using these independent variables and their pairwise interactions, including the complete model and a null model. Predictors were standardized (mean =0, SD = 0.5) prior to the analysis in order to simplify interpretation of model averaged parameter estimates, as recommended by Grueber et al. (2011). Akaike information criterion corrected for low values of the ratio between sample size and parameters (i.e., AICc) were used to evaluate each model within the set of candidate models (Burnham & Anderson, 2002). Candidate models were then ranked from best to worst based on their deltaAICc values. Akaike weights and evidence ratios were used to assess the relative strengths of each candidate model.

Because we were interested in determining which factors have the strongest effect on foraging success during a trial, we performed a model averaging procedure on the 95% candidate set of models (i.e., top models with a cumulative weight of 95%) using a full averaging method (i.e., ‘zero method’) implemented in the ‘MuMIn’ package (Burnham & Anderson, 2002; Nakagawa & Freckleton, 2011). The relative importance (RI) of the parameters was calculated by summing the Akaike weights from the models in which they appear, using the 95% candidate set of models. RI values close to 1 indicate that the parameter appears in the best supported models, while RI values close to 0 indicate that the parameters appear in the least supported models.

We performed the same AICc model averaging procedure for each day separately in order to investigate whether there were changes in the relative importance of the predictors on foraging success as individuals acquire more reliable private information (i.e., over repeated days of testing) and because ‘day’ had a strong effect on most of the variables of interest. Doing so also maintained parsimony in the model construction.

All data were analysed using the ‘lme4’ (Bates et al., 2015), ‘MASS’ (Venables & Ripley, 2002), ‘car’ (Fox & Weisberg, 2019) and ‘MuMIn’ (Barton, 2020) packages for modelling and visualized using the ‘ggplot2’ (Wickham, 2016) package in R 3.6.2. (R Core Team, 2019).

## RESULTS

### Depletion time

There was a significant effect of the interaction between treatment and day on the depletion time measured in shoals foraging on aggregated and scattered treatments (negative binomial GLMM: χ2 = 6.26, df = 1, *p* = 0.012). Depletion time decreased over the first four days of the experiment, before reaching a plateau at around 60 seconds on the fifth day of testing (Figure 2).

### Movement characteristics

There was a significant effect of the interaction between treatment and day on the total distance travelled (GLMM, Day: Treatment: χ2 = 104.89, df = 6, *p* < 0.001). Distances travelled increased over the first three days for individuals foraging on either aggregated or scattered prey distribution, whereas the distances travelled by individuals in the control treatment decreased over the first five tests (Figure 3A).

**Figure 3.**
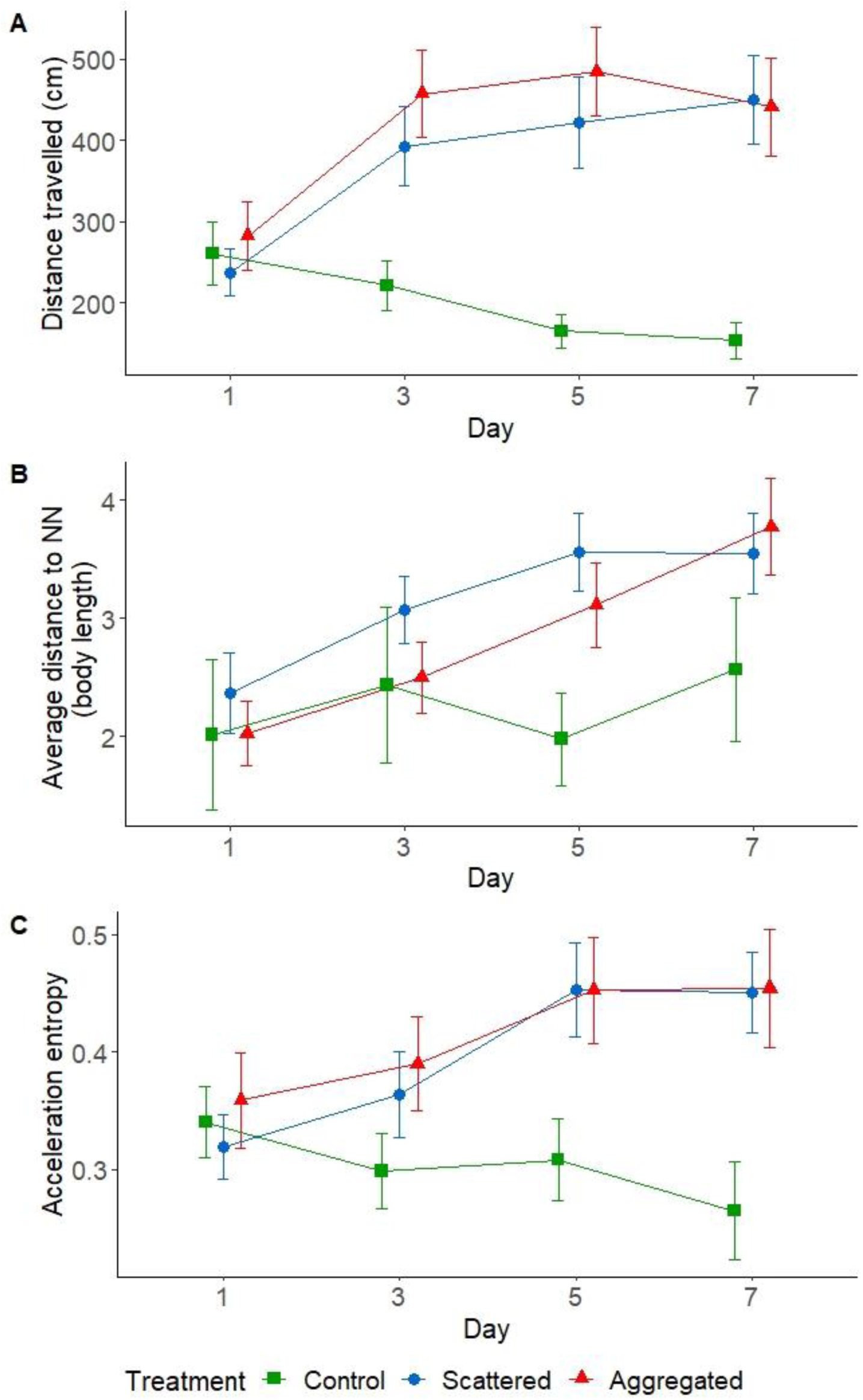
Temporal changes of individuals’ movement characteristics as a function of treatments (mean ± 95%CI). (A) Total distance travelled (in cm), calculated as the cumulated value over all frames spent moving (i.e., speed greater than 0.5 BL.s^-1^). (B) Average distance to nearest neighbour (in body length), calculated as the median value over all frames spent moving. (C) Acceleration entropy calculated as the median of entropy values of all bouts of at least 15 consecutive frames spent moving.

There was a significant effect of the interaction between treatment and day on the average distance to the nearest neighbour (GLMM, Day: Treatment: χ2 = 28.72, df = 6, *p* < 0.001). Distances to nearest neighbour tended to remain consistent over the repeated tests in fish from the control treatment, whereas individuals from both aggregated and scattered treatments increased their distance to their nearest neighbour over at least the first five repetitions of the foraging task (Figure 3B).

There was a significant effect of the interaction between treatment and day on the acceleration entropy (GLMM, Day: Treatment: χ2 = 56.68, df = 6, *p* < 0.001). Acceleration entropy increased over the first 5 days of testing before stabilising in fish from both aggregated and scattered treatments, whereas it slightly decreased over the repeated tests in fish from the control treatment (Figure 3C).

#### Foraging success

Distance to NN, treatment and their interaction, were identified from the model averaging procedure to be the most important predictors of foraging success on day 1 (Table 1). These terms occurred in the best three models of the 95% candidate set of models (models 32, 22 and 64; Table S2). The second and third best supported models both had a strong support (i.e., delta AICc < 4) and evidence ratios smaller than 3.1 (Table S2). Individuals that maintained greater distances between themselves and their nearest neighbour had a greater foraging success when the prey were aggregated, whereas greater foraging success decreased when individuals maintained greater distances between themselves and their nearest neighbour when the prey were scattered in the arena (Figure 4A). Note that distance travelled and the interaction between treatment and distance travelled also had a strong relative importance (RI = 0.77 for both) despite being statistically non-significant (Table 1).

**Table 1.** Standardized model averaged coefficient estimates from the 95% candidate set of models to determine the best predictors of foraging success within each day of testing. Occurrence denotes the number of models that contain the term. Standardized model averaged coefficient estimates (based on the 95% candidate set of models) are reported with the standard error (SE) and the 95% confidence interval (95%CI). The relative importance (RI) of each term is the sum of the Akaike weights over the models in which that tern appears (based on the 95% candidate set of models). Important terms of significant effects are indicated in bold within each distinct day, and marginally non-significant effects are indicated by *.

| Day | Terms | Occurrence | Estimates | SE | 95%CI | RI |
| --- | --- | --- | --- | --- | --- | --- |
| 1 | Intercept | 3 | 0.983 | 0.116 | [0.756, 1.21] | - |
| 1 | Distance travelled | 2 | -0.029 | 0.162 | [-0.346, 0.289] | 0.77 |
| 1 | **Distance to NN** | **3** | **0.305** | **0.15** | **[0.012, 0.599]** | **1** |
| 1 | **Treatment** | **3** | **-0.16** | **0.227** | **[-0.604, 0.284]** | **1** |
| 1 | Distance travelled x Distance to NN | 1 | 0.022 | 0.172 | [-0.315, 0.359] | 0.187 |
| 1 | Distance travelled x Treatment | 2 | -0.676 | 0.489 | [-1.634, 0.282] | 0.77 |
| 1 | **Distance to NN x Treatment** | **3** | **1.011** | **0.296** | **[0.43, 1.592]** | **1** |
| 3 | Intercept | 7 | 1.619 | 0.047 | [1.528, 1.71] | - |
| 3 | **Distance travelled** | **7** | **0.521** | **0.089** | **[0.347, 0.694]** | **1** |
| 3 | **Distance to NN** | **7** | **0.297** | **0.093** | **[0.115, 0.479]** | **1** |
| 3 | Treatment | 5 | -0.031 | 0.071 | [-0.17, 0.108] | 0.423 |
| 3 | Distance travelled x Distance to NN | 3 | 0.036 | 0.127 | [-0.213, 0.284] | 0.265 |
| 3 | Distance travelled x Treatment | 2 | 0.01 | 0.064 | [-0.116, 0.136] | 0.098 |
| 3 | Distance to NN x Treatment | 1 | -0.006 | 0.053 | [-0.11, 0.097] | 0.069 |
| 5 | Intercept | 8 | 1.708 | 0.044 | [1.621, 1.795] | - |
| 5 | **Distance travelled** | **8** | **0.399** | **0.088** | **[0.226, 0.573]** | **1** |
| 5 | Distance to NN | 5 | 0.004 | 0.058 | [-0.11, 0.117] | 0.322 |
| 5 | Treatment | 5 | -0.011 | 0.053 | [-0.114, 0.093] | 0.334 |
| 5 | Distance travelled x Distance to NN | 2 | 0.007 | 0.063 | [-0.116, 0.13] | 0.076 |
| 5 | Distance travelled x Treatment | 1 | -0.003 | 0.044 | [-0.089, 0.082] | 0.058 |
| 5 | Distance to NN x Treatment | 1 | -0.009 | 0.057 | [-0.12, 0.103] | 0.038 |
| 7 | Intercept | 11 | 1.738 | 0.044 | [1.65, 1.825] | - |
| 7 | Distance travelled | 6 | 0.096 | 0.102 | [-0.103, 0.296] | 0.632 |
| 7 | Distance to NN | 6 | -0.011 | 0.055 | [-0.12, 0.097] | 0.301 |
| 7 | Treatment | 6 | -0.003 | 0.048 | [-0.098, 0.092] | 0.295 |
| 7 | Distance travelled x Distance to NN | 1 | -0.002 | 0.043 | [-0.086, 0.082] | 0.036 |
| 7 | Distance travelled x Treatment | 1 | 0.007 | 0.05 | [-0.091, 0.105] | 0.047 |
| 7 | Distance to NN x Treatment | 1 | -0.005 | 0.045 | [-0.094, 0.084] | 0.021 |

**Figure 4.**
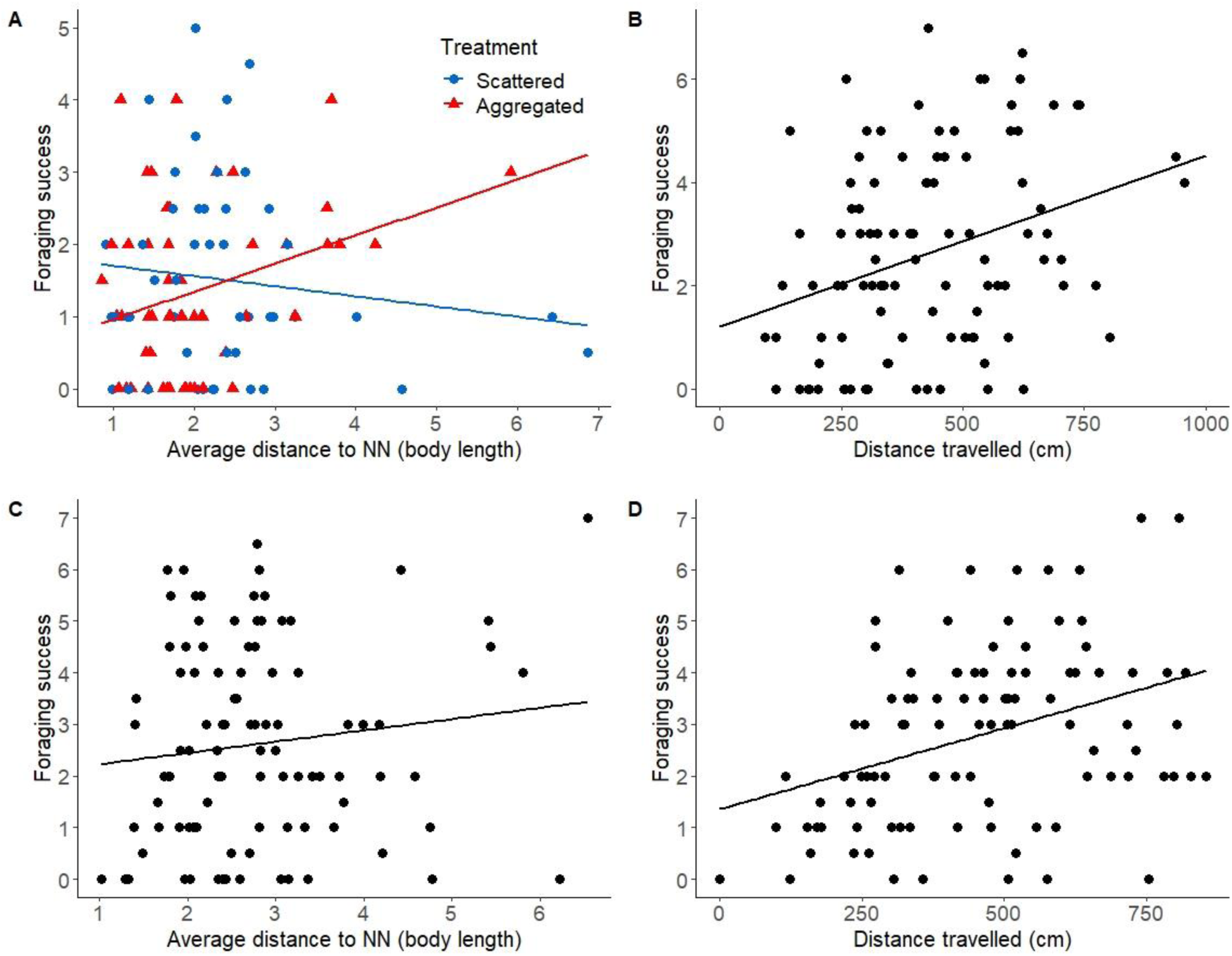
Visualisation of the effects of the movement characteristics identified to be significant predictors of foraging success from the model averaging procedure performed within days 1, 3 and 5. (A) The effect of the interaction between the total distance travelled (in cm) and treatment on individual foraging success on day 1. (B) The effect of total distance travelled (in cm) on individual foraging success on day 3. (C) The effect of the average distance to nearest neighbour (in average body length unit) on individual foraging success on day 5. (D) The effect of total distance travelled (in cm) on individual foraging success on day 3.

Distance to NN and the distance travelled were identified from the model averaging procedure to be the most important predictors of foraging success on day 3 (Table 1). Both terms occurred in the best three models of the 95% candidate set of models (models 7, 8 and 39; Table S2). The second and third best supported models both had a substantial support (i.e., delta AICc < 2) and evidence ratios smaller than 2.5 (Table S2). Foraging success increased with distance travelled (Figure 4B) and distance to NN (Figure 4C) independently of the treatment.

Distance travelled was identified from the model averaging procedure to be the most important predictor of foraging success on day 5 (Table 1). This term occurred in the best three models of the 95% candidate set of models (models 3,4 and 7; Table S2). The second and third best supported models both had a strong support (i.e., delta AICc < 4) and evidence ratios smaller than 3 (Table S2). Individuals that travelled greater distances had greater foraging success, independently of the treatment (Figure 4D).

On day 7, distance travelled was identified from the model averaging procedure to be the most important predictor of foraging success on day 7, with a relative importance of 0.63, but had a very small effect which did not statistically differ from zero (Table 1).

## DISCUSSION

We showed that the movement characteristics of individuals changed over repeated foraging trials. These changes mostly occurred within the first 3 to 5 days of our repeated tests and differed between control and foraging treatments (i.e., scattered and aggregated treatments). Unlike control individuals, fish foraging on scattered and aggregated prey showed changes in all three movement variables considered in this study, and demonstrated a significant increase in their collective foraging efficiency over the first 3 days of testing. Overall, we found pronounced increases in distance travelled between days 1 and 3 as well as in distance to nearest neighbour between days 1 and 5 in fish from scattered and aggregated treatments, which contrasted with control fish that maintained consistent values of distance travelled and distance to NN over similar times. Acceleration entropy also increased in foraging fish over the first 5 days, with the most pronounced increase occurring between days 3 and 5, which differs from control fish whose entropy tended to remain unchanged over time. As hypothesised, all three variables seemed to reach a plateau initiated somewhere between days 3 and 5. We found no effect of the spatial distribution of prey on the movement of foraging individuals. Furthermore, we found that the relative importance of the variables considered here changed over time; with an important effect of social proximity on the testing days 1 and 3, while search effort had an important effect on variation in foraging success on tests 3 and 5.

The changes occurring over time suggest an important effect of the knowledge individuals possess about their foraging context on their movement characteristics. Without any prior information on the surrounding environment (i.e., day 1), individuals from all treatments travelled similar distances, were equally unpredictable in their variation in speed, and positioned themselves within approximately 2 body lengths from their nearest neighbour. Such relatively close proximity to the nearest neighbour is typically observed in species known for their strong schooling behaviour when exploring unfamiliar experimental or natural environments (e.g., Calovi et al., 2018; Schaerf et al., 2017; Viscido et al., 2004; Ward et al., 2017). As hypothesised, our results suggest that individuals displayed grouping behaviour in absence of prior information, by maintaining close social proximity. This was also supported by investigations performed at the group level, showing that individuals tended to leave or join others on relatively few occasions (see Supporting Information). Social proximity is often assumed to enhance integration of information, particularly when exploring novel environments (Berdahl et al., 2013; Couzin, 2007; Sumpter et al., 2008). In the absence of prior information, grouping behaviour in fish has been found to enhance both individual foraging and the ability to detect and avoid potential predators (Pitcher et al., 1982; Ward et al., 2011). As hypothesised, social proximity was the most important predictor of foraging success during the first foraging test, and depended on the spatial distribution of prey. Although we found that grouping behaviour was most pronounced on that day, individuals that maintained greater distances to their nearest neighbour tended to have a higher foraging success when prey were aggregated, but a lower foraging success when prey were scattered. This result suggests that adopting a more asocial and risk-taking strategy by staying further away from conspecifics may confer a greater payoff only when successful discoveries lead to larger intake opportunities, as was the case in the aggregated treatment. Alternatively, individuals foraging on scattered prey may maximize their foraging success by adopting a social strategy. It may be possible that foragers increased their feeding opportunity by maintaining a close proximity to their conspecifics which would make it easier to kleptoparasitise successful neighbours (Hansen et al., 2016).

Repeated testing offered all individuals the opportunity to reinforce their private information, which included the presence (or not, in the case of the control shoals) of food in the arena, its spatial occurrence, and the presence of a consistent number of competitors. This is likely to have driven the changes observed in the distance travelled, the overall social proximity and unpredictability in acceleration. The high relative importance of social proximity in explaining variation in foraging success on day 3 suggests that maintaining longer distances from competitors is of primary importance. An individual which just discovered a prey and is in the process of consuming a prey provides social information which attracts conspecifics (Harpaz & Schneidman, 2020). Monitoring positions of local neighbours at any time and maintain appropriate distances (e.g., by constantly avoiding swimming towards conspecifics and fleeing away from those getting too close) may prevent the loss of a prey to larger competitors. This mechanism may have led to increasing the total distance travelled, and ultimately resulted in a greater foraging success for individuals that remained away from any conspecific on day 3. Such interactions based on social information and driven by competition would explain the changes in acceleration entropy observed in scattered and aggregated treatments from the third day onwards. Our additional investigations performed at the group level support these explanations by demonstrating that grouping events became more frequent but tended to be short-lived from day 3 onwards (see Supporting Information).

As the majority (if not all) of the individuals have integrated similar information after a certain number of repetitions of the foraging task, simply adjusting distance to neighbours may no longer be enough to score a high foraging success. The fact the distance travelled was the most important predictor of foraging success on days 3,5 and 7 (despite non statistical significance) suggests that feeding performance may simply come down to being able to sample the greatest proportion of the arena possible in a short time (resulting in travelling greater distances) when all foragers possess information about food spatial occurrence. It is possible that when information is widely spread within the group, individuals may not get much benefit out of it, and therefore rely mostly on their search effort. Our data demonstrates that this transition occurred after day 3, suggesting that individuals switch from paying attention to their social environment and responding to it accordingly in the absence of reliable information, to prioritizing search effort to cope with competition coming from other informed foragers. Although this transition occurred on the same day the collective foraging efficiency reached its maximum, establishing the relation of causality between such behavioural switch and the foraging efficiency remains to be addressed in future investigations.

Our results show evidence that while the presence of food affects individual movement throughout the repeated tests, there was no effect of the spatial distribution of prey. It is known that fish can acquire a variety of information about their current ecological context by sensing olfactory cues diluted in the water, which can lead to behavioural changes (e.g., Hoare et al., 2004; Ferrari et al., 2010; Sosna et al. 2019). Previous work on schooling fish suggested that individuals are less frequently observed at close proximities form one another and have higher magnitude of acceleration in the presence of food odour (Schaerf et al., 2017), which is in line with our findings. Furthermore, fish can retain information about temporal and spatial food occurrence and use this to improve foraging (Reebs, 2000; Van Bergen et al., 2004). Yet, our results show no difference in changes of movement characteristics between fish from the aggregated and scattered treatments, which suggest that knowledge of prey availability triggers behavioural responses that are independent of their spatial distribution. This surprising result goes against our initial working hypothesis and differs from previous work on juvenile walleye pollock (*Theragra chalcogramma)* conducted by Ryer & Olla (1995), as well as a study on a broad range of open-ocean predatory fish (Humphries et al., 2010), but is in line with a study on foraging groups of three-spined sticklebacks (*Gasterosteus aculeatus*) conducted by Hansen et al. (2016). Although our arena is consistent in size with those used in prior studies, it is possible that the distance between prey items in the scattered treatment did not differ sufficiently from the distance separating each pair of patches used in the aggregated treatment. When foraging in the wild, individuals may have to explore larger portions of the environment between successful samplings, inducing higher energetic and time costs from search and transport. Such costs may be compensated by moving in a manner that maximizes food discovery and reduces costs of competition (e.g., grouping when foraging on clumped patches of food, foraging asocially when the food is distributed across the environment). Additionally, the temporal uncertainty of food occurrence in natural environments may reinforce the need for pronounced behavioural adjustment in response to different spatial distribution of resources. Therefore, the scale used to test the two distributions of prey in our study, combined with the experimental control for temporal uncertainty by making food available within each daily test, may not have imposed high enough costs of transport and search required to observe distinct movement patterns.

Overall, we found that behavioural changes appeared to be driven by individual information about the foraging context, including learned presence of food and competitors, but were unaffected by the spatial distribution of that food. As individuals reinforced their information about foraging conditions, they may have to adjust their behavioural strategy to cope with competitors by switching from maintaining an adequate social proximity to maximizing search effort. Further research considering temporal unpredictability and a larger experimental scale, where scattered (or patches of) prey would be separated by greater distances, would likely promote the use of fundamentally different movement strategies at the individual level and would ultimately provide a better understanding of how resource-driven individual movement strategies affect group self-organisation.

## ACKNOWLEDGEMENTS

We thank Lysanne Snijders and Darren Croft for constructive comments on the first version of the manuscript. This work was supported by a Barry Jonassen Award granted by the Australian Society for Fish Biology to G.P.F.M. and an ARC Discovery Project Grant DP190100660 to A.J.W.W.

## Notes

**CONFLICT OF INTEREST** The authors declare no competing interests.

### Competing Interest Statement

The authors have declared no competing interest.

### Summary of Updates

Results were revised. Main text was updated.

