## Supplementary Material for "Changes in movement characteristics in response to private and social information acquisition of socially foraging fish"

**1. BODY SIZE**

A one-way ANOVA was conducted to examine whether the groups average body size different across treatments. There was no statistically significant difference across treatments (F_2,32_ = 0.073, p = 0.93).

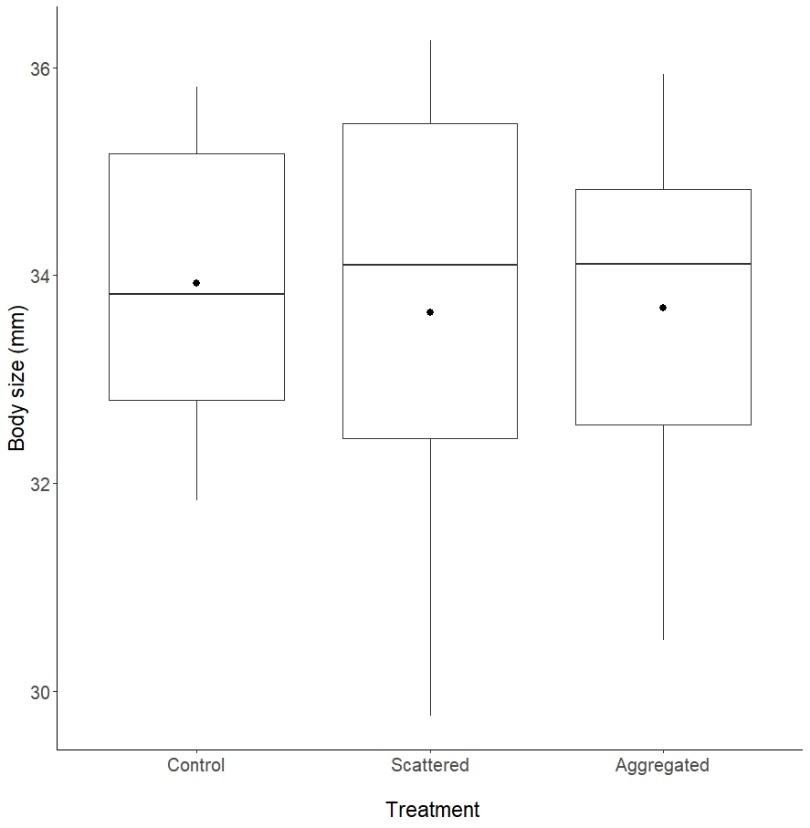

**Figure S1.** Within groups average body size across treatments (thick line = median, black dot = mean).

**2. MODEL AVERAGING**

**2.1 Collinearity between the continuous variables of interest**

Collinearity between continuous variables was examined using correlation tests prior to performing the model averaging. Correlations were estimated using the Kendall’s rank order correlation coefficient (tau) rather that Spearman’s rho, as it produces more reliable confidence intervals in the absence of ties in the data (Puth et al., 2015).

**Table S1.** Correlations between the continuous variables of interest within each day. Kendall correlation coefficients (*tau*) are presented with 95% confidence intervals. Significant correlation coefficients after correction for multiple comparisons (following Benjamini-Hochberg procedure) are indicated in bold.

| Day | Variable_1 | Variable_2 | Kentall tau | 95%CI |
| --- | --- | --- | --- | --- |
| 1 | Distance travelled | Distance to NN | **0.15** | [-0.02, 0.31] |
| 1 | Distance travelled | Acceleration entropy | **0.48** | [0.34, 0.60] |
| 1 | Distance to NN | Acceleration entropy | **0.18** | [0.01, 0.35] |
| 3 | Distance travelled | Distance to NN | 0.01 | [-0.16, 0.18] |
| 3 | Distance travelled | Acceleration entropy | **0.40** | [0.25, 0.53] |
| 3 | Distance to NN | Acceleration entropy | -0.02 | [-0.19, 0.15] |
| 5 | Distance travelled | Distance to NN | **0.13** | [-0.04, 0.29] |
| 5 | Distance travelled | Acceleration entropy | **0.45** | [0.31, 0.57] |
| 5 | Distance to NN | Acceleration entropy | **0.17** | [0, 0.32] |
| 7 | Distance travelled | Distance to NN | **0.15** | [-0.02, 0.31] |
| 7 | Distance travelled | Acceleration entropy | **0.48** | [0.34, 0.60] |
| 7 | Distance to NN | Acceleration entropy | **0.18** | [0.01, 0.35] |

**2.2 Set of candidate models used for model comparisons**

**Table S2.** Set of candidate models based on the 95% cumulative weights criterion used for model comparisons based on Akaike’s Information Criterion. Generalized linear mixed-effects models explain foraging success by changes in treatment (T), distance to nearest neighbour (NND), distance travelled (D) and their interactions. All models include shoal identity as random term. All compared models are ranked within each day based on decreasing delta AIC. (K= number of fitted parameters, logLik = log likelihood, AICc = corrected Akaike’s Information Criterion, ΔAICc = delta AICc, w*_i_* = model Akaike weights, ER = evidence ratio**).**

| Day | Model | Terms | K | logLik | AICc | ΔAICc | w*_i_* | Rank | ER |
| --- | --- | --- | --- | --- | --- | --- | --- | --- | --- |
| 1 | 32 | T + D + NND + T:D + T:NND | 7 | -218.718 | 452.654 | 0 | 0.51 | 1 | 1 |
| 1 | 22 | T + NND + T:NND | 5 | -221.937 | 454.513 | 1.859 | 0.201 | 2 | 2.533 |
| 1 | 64 | T + D + NND + T:D + T:NND + D:NND | 8 | -218.669 | 454.921 | 2.267 | 0.164 | 3 | 3.107 |
| 3 | 7 | D + NND | 4 | -299.664 | 607.748 | 0 | 0.383 | 1 | 1 |
| 3 | 8 | T + D + NND | 5 | -299.337 | 609.313 | 1.564 | 0.175 | 2 | 2.186 |
| 3 | 39 | D + NND + D:NND | 5 | -299.449 | 609.537 | 1.789 | 0.156 | 3 | 2.446 |
| 3 | 40 | T + D + NND + D:NND | 6 | -299.198 | 611.299 | 3.551 | 0.065 | 4 | 5.903 |
| 3 | 24 | T + D + NND + T:NND | 6 | -299.199 | 611.301 | 3.553 | 0.065 | 5 | 5.909 |
| 3 | 16 | T + D + NND + T:D | 6 | -299.207 | 611.316 | 3.568 | 0.064 | 6 | 5.954 |
| 3 | 48 | T + D + NND + T:D + D:NND | 7 | -298.921 | 613.06 | 5.311 | 0.027 | 7 | 14.232 |
| 5 | 3 | D | 3 | -264.521 | 535.295 | 0 | 0.429 | 1 | 1 |
| 5 | 4 | T + D | 4 | -264.448 | 537.322 | 2.027 | 0.156 | 2 | 2.755 |
| 5 | 7 | D + NND | 4 | -264.517 | 537.46 | 2.165 | 0.145 | 3 | 2.952 |
| 5 | 12 | T + D + T:D | 5 | -264.39 | 539.425 | 4.13 | 0.054 | 4 | 7.885 |
| 5 | 39 | D + NND +D:NND | 5 | -264.407 | 539.459 | 4.164 | 0.053 | 5 | 8.020 |
| 5 | 8 | T + D + NND | 5 | -264.448 | 539.541 | 4.246 | 0.051 | 6 | 8.356 |
| 5 | 24 | T + D + NND + T:NND | 6 | -263.683 | 540.279 | 4.984 | 0.035 | 7 | 12.085 |
| 5 | 40 | T + D + NND + D:NND | 6 | -264.369 | 541.651 | 6.355 | 0.018 | 8 | 23.987 |
| 7 | 3 | D | 3 | -222.958 | 452.189 | 0 | 0.287 | 1 | 1 |
| 7 | 1 | 1 | 2 | -224.558 | 453.25 | 1.061 | 0.169 | 2 | 1.700 |
| 7 | 7 | D + NND | 4 | -222.93 | 454.321 | 2.132 | 0.099 | 3 | 2.904 |
| 7 | 4 | T + D | 4 | -222.954 | 454.368 | 2.179 | 0.097 | 4 | 2.973 |
| 7 | 5 | NND | 3 | -224.323 | 454.919 | 2.73 | 0.073 | 5 | 3.916 |
| 7 | 2 | T | 3 | -224.55 | 455.373 | 3.184 | 0.058 | 6 | 4.914 |
| 7 | 12 | T + D + T:D | 5 | -222.609 | 455.915 | 3.726 | 0.045 | 7 | 6.443 |
| 39 | 39 | D + NND + D:NND | 5 | -222.896 | 456.489 | 4.3 | 0.033 | 8 | 8.585 |
| 8 | 8 | T + D + NND | 5 | -222.928 | 456.553 | 4.364 | 0.032 | 9 | 8.864 |
| 6 | 6 | T + NND | 4 | -224.321 | 457.102 | 4.913 | 0.025 | 10 | 11.664 |
| 22 | 22 | T + NND + T:NND | 5 | -223.409 | 457.516 | 5.327 | 0.02 | 11 | 14.346 |

**2. Investigations of grouping behaviour**

**2.1 Hypothesis**

We hypothesized that the individual searching strategy favoured with each type of prey distribution and/or the reliability of provide information (i.e., day) would results in different patterns of self-organisation at the level of the group. Considering that intraspecific competition may act as a driving force jointly with prey distribution, we predicted that the tested shoals would be more inclined to display a cohesive and collective search strategy in the absence of information (i.e., on day 1) and when the discovery of prey is less frequent (i.e., harder to locate due to lower occurrence) but highly rewarding (i.e., aggregated treatment), while those foraging on more evenly distributed prey associated with higher levels of scramble competition would display a ‘solitary’ type of searching strategy and would distribute themselves across the arena.

**2.2 Identification of grouping behaviour**

Using the tracking data, we calculated the Euclidean distance between each pair of fish at each time step. We considered that two individuals were grouping if they were within 4 body lengths apart from each other (from now referred to as ‘neighbours’) - a threshold commonly used to determine group membership in fish shoals (Pitcher et al., 1983; Hoare et al., 2004; Herbert-Read et al., 2017) - and that an individual could join an existing group by positioning itself within 4 body lengths from at least one group member. Using these criteria, we determined whether each fish was alone or part of a group, and we quantified their number of neighbours (referring to the number of fish within a radius of 4 body lengths) at each time step. Based on the number of fish that were identified to be part of a group, the sum of their number of neighbours, and the number of individuals that had only one neighbour, grouping behaviour occurring within our 4-fish shoals could fall into one of eleven different grouping patterns (Table S3). From these data, we then recorded the type of grouping structure that occurred at each time step and extracted the identities of the individuals in the group. Finally, we defined a grouping event as a sequence of successive time steps for which there were no changes in group members identities but could contain changes in grouping structure within a given group size (e.g., transitioning from structure 4 to 5, Table S3). For any grouping event that lasted longer than 1 frame, we recorded its length (number of frames) and the size of the group (e.g., group size of 2 indicating a grouping event occurring between two fish, etc.).

**2.3 Analysis of grouping behaviour**

We analysed the influence of treatment and days on the individual proportion of time spent grouping (i.e., identified to be part of grouping events, parsed as a data frame) using a GLMM fitted with a binomial error distribution and the logit link function. The dependent variable consisted of a two-column vector response variable, by binding together the number of frames spent grouping and the number of frames spent not grouping. Fish identity was nested within the shoal identity as random effects.

We analysed the influence of treatment, day and group size on the number of grouping events recorded within the 60 seconds of observation using a Generalized Linear Mixed Effects model (GLMM) fitted with a negative binomial error distribution (due to an overdispersion of the data but without zero inflation), using the ‘glmmTMB’ package (Brooks et al., 2017). The fixed effects used in the model included day, treatment, group size and all their pairwise interactions. Shoal identity was used as a random effect.

We analysed the influence of treatment, day and group size on the duration of the recorded grouping events using a GLMM fitted with a negative binomial error distribution. The fixed effects used in the model included day, treatment, group size and all their pairwise interactions. Shoal identity was used as a random effect.

All models were first simplified by removing non-significant interactions, by assessing the AIC value of models with and without the tested terms. The significance of the terms kept in the reduced models were assessed using either type II or type III ANOVAs.

**Table S3.** Different grouping structures observable in a group of 4 fish, using a given thresholding distance (4 average body lengths) as grouping criteria.

| **Description** | **Group size** | **Grouping pattern** |
| --- | --- | --- |
| (1) All individuals are further than 4 body lengths away from each other (i.e., no grouping). | 0 |  |
| (2) Only two individuals are within 4 body lengths away from each other, two individuals are alone. | 2 |  |
| (3) All individuals are within 4 body lengths away from a single neighbour, forming two pairs. | 2 |  |
| (4) Two individuals are within 4 body lengths away from the same individual but are further away from each other, and one individual alone. | 3 |  |
| (5) Tree individuals are within 4 body lengths away from each other, and one individual is alone. | 3 |  |
| (6) Two individuals are within 4 body lengths from two different individuals which are themselves within 4 body lengths away from each other. | 4 |  |
| (7) Three individuals are within 4 body lengths from a common neighbour. | 4 |  |
| (8) All four individuals are within 4 body lengths away from two other individuals. | 4 |  |
| (9) Three individuals are within 4 body lengths away from each other, and one individual is within 4 body lengths from one of them. | 4 |  |
| (10) Three individuals are within 4 body lengths away from each other, and one individual is within 4 body lengths from two of them. | 4 |  |
| (11) All four individuals are within 4 body lengths away from each other. | 4 | 4 (or less) body lengths |

**2.4 Results**

There was a significant effect of the interaction between treatment and day on the proportion of time that individuals spent grouping (Table S4). Time spent grouping was greater for individuals from the control treatment and remained constant over days, whereas individuals from scattered and aggregated treatments decreased their time spent grouping over days (Figure S2A).

There were significant effects of the interactions between treatment and day, the interaction between treatment and group size, and the interaction between day and group size on the number of grouping events (Table S4). The number of grouping events recorded in the control group remained consistent across days, whereas it increased between the first and third day in the scattered and aggregated treatments. Grouping events recorded from day 3 onwards in the scattered and aggregated treatments were nearly twice as many as those recorded in the control treatment (Figure S2B). Shoals from the scattered and aggregated treatments displayed three times more grouping events of 2 than control shoals, and twice as many grouping events of 3 as control shoals. The number of grouping events of 4 did not differ between treatments (Figure S2C). The number of grouping events of 4 was consistently lower than the number of grouping events of 3 across days, which were themselves significantly lower than the number of grouping events of 2 for all days except for day 1 (Figure S2D).

There were significant effects of the interaction between treatment and group size, the interaction between treatment and group size, and the interaction between day and group size on the duration of grouping events (Table S4). The average duration of grouping events in the control group remained consistent across days, whereas it decreased between the first and third day in the scattered and aggregated treatments. Grouping events recorded from day 3 onwards in the scattered and aggregated treatments were more than twice shorter than those recorded in the control treatment (Figure S2E). Grouping events occurring in shoals from the scattered and aggregated treatments did not differ in duration within each group size, but were significantly lower than those occurring in control shoals independently of the group size (Figure S2F). Grouping events of 4 lasted significantly longer than groups of 2 and 3, which tended to have similar durations, on all days except for day 7 where grouping events of all sizes had equal durations (Figure S2G).

**Table S4.** Results of the modelling investigations performed on the grouping behaviour. Significant effects are indicated in bold characters.

| Dependent variable | Modelling function (family) | Terms | Chi-square (χ2) | DF | *P* |
| --- | --- | --- | --- | --- | --- |
| Proportion of time grouping | glmer (binomial (link="logit")) | **Treatment:Day** | **4697.487** | **6** | **<0.001** |
|  |  | **Treatment** | **50.302** | **2** | **<0.001** |
|  |  | **Day** | **20656.184** | **3** | **<0.001** |
| Number of grouping events | glmmTMB (nbinom) | **Treatment:Day** | **14.882** | **6** | **0.021** |
|  |  | **Treatment:Group size** | **56.780** | **4** | **<0.001** |
|  |  | **Day:Group size** | **33.148** | **6** | **<0.001** |
|  |  | **Treatment** | **30.945** | **2** | **<0.001** |
|  |  | **Day** | **13.211** | **3** | **0.004** |
|  |  | **Group size** | **236.149** | **2** | **<0.001** |
| Duration of grouping events | glmer.nb() | **Treatment:Day** | **55.222** | **6** | **<0.001** |
|  |  | **Treatment:Group Size** | **29.216** | **4** | **<0.001** |
|  |  | **Day:Group Size** | **37.855** | **6** | **<0.001** |
|  |  | **Treatment** | **59.308** | **2** | **<0.001** |
|  |  | **Day** | **64.224** | **3** | **<0.001** |
|  |  | **Group Size** | **33.628** | **2** | **<0.001** |

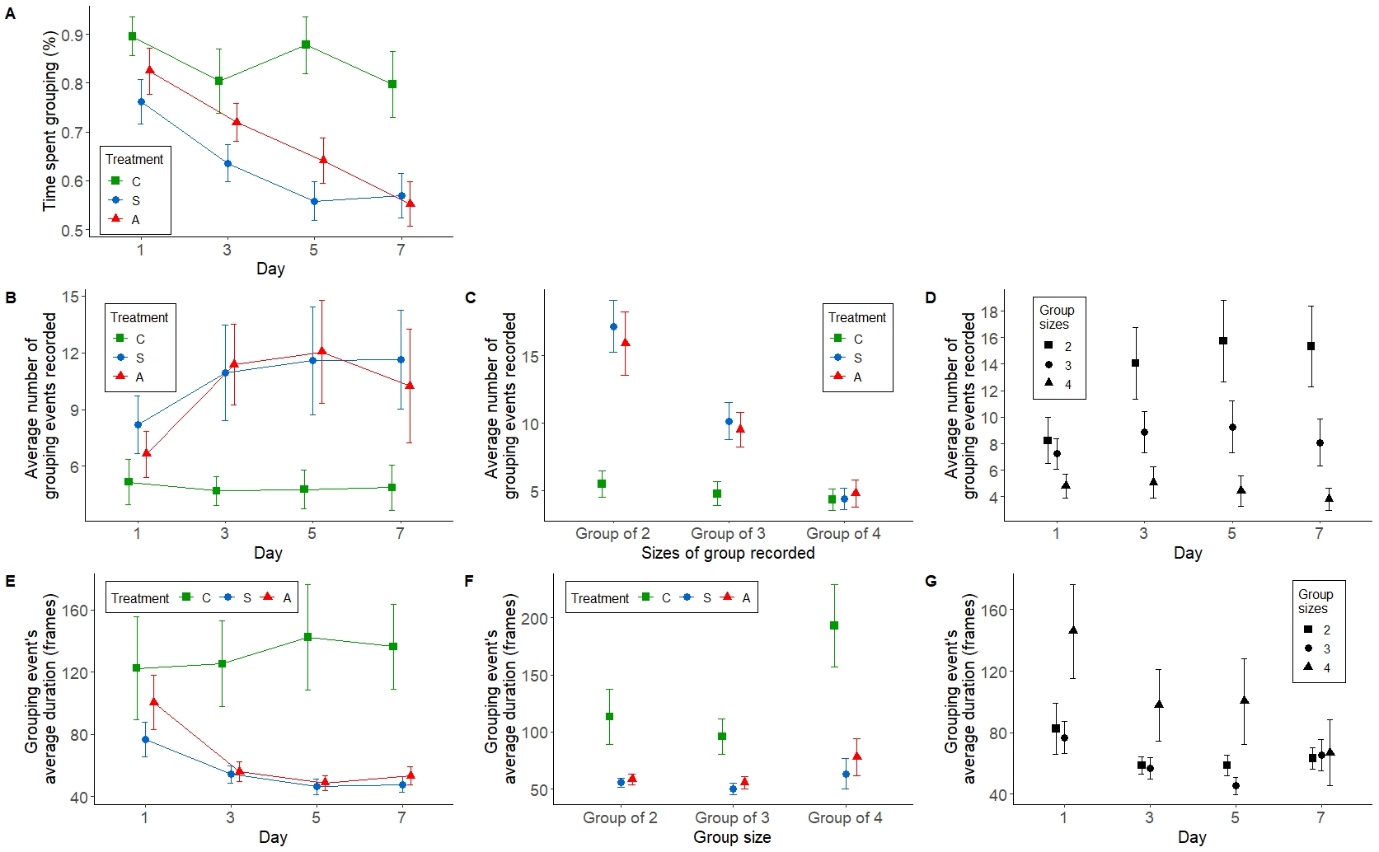

**Figure S2.** Visualization of the analyses of grouping behaviour. (A) The effects of the interaction between days and treatment on the individual time spent grouping with other conspecifics (mean ± 95%CI). (B) The effects of the interaction between days and treatment on the number of grouping events recorded (mean ± 95%CI). (C) The effects of the interaction between group size and treatment on the duration of grouping events (mean ± 95%CI). (D) The effects of the interaction between group size and days on the duration of grouping events (mean ± 95%CI). (E) The effects of the interaction between days and treatment on the duration of grouping events (mean ± 95%CI). (F) The effects of the interaction between group size and treatment on the duration of events (mean ± 95%CI). (G) The effects of the interaction between group size and days on the duration of events (mean ± 95%CI).
